## Supplementary figures for "Impact of a candidate vaccine on the dynamics of salmon lice (*Lepeophtheirus salmonis*) infestation and immune response in Atlantic salmon (*Salmo salar* L.)": S2_Fig.pdf

### PCA analysis- Head kidney

A

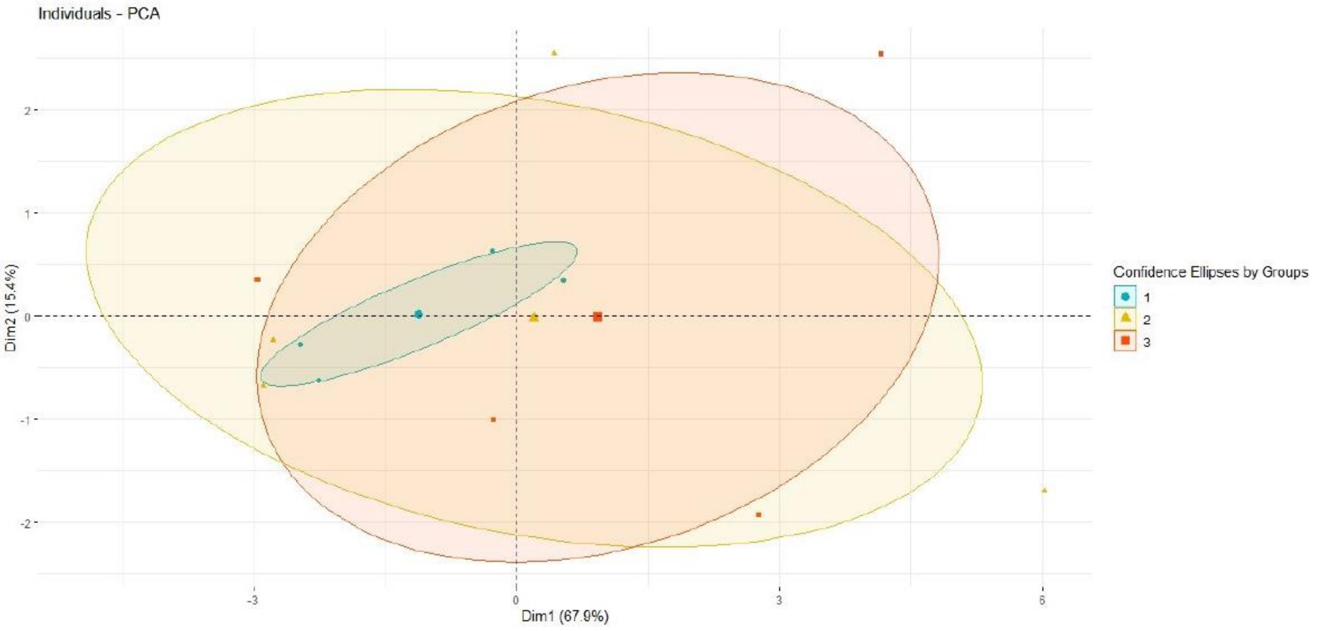


B

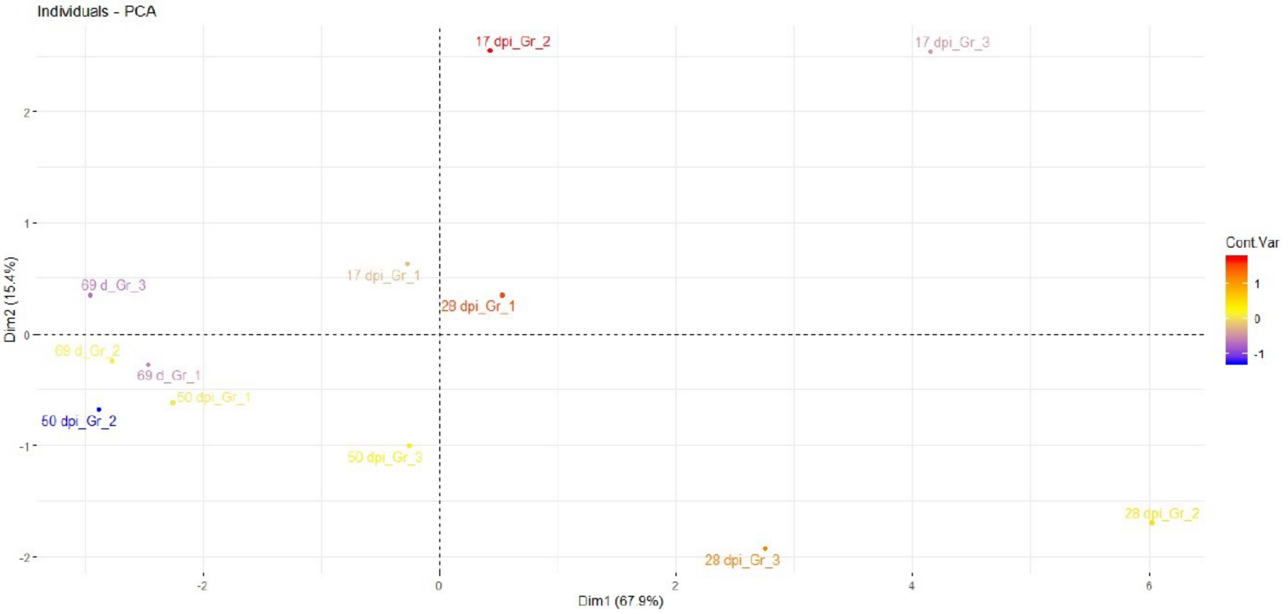


C

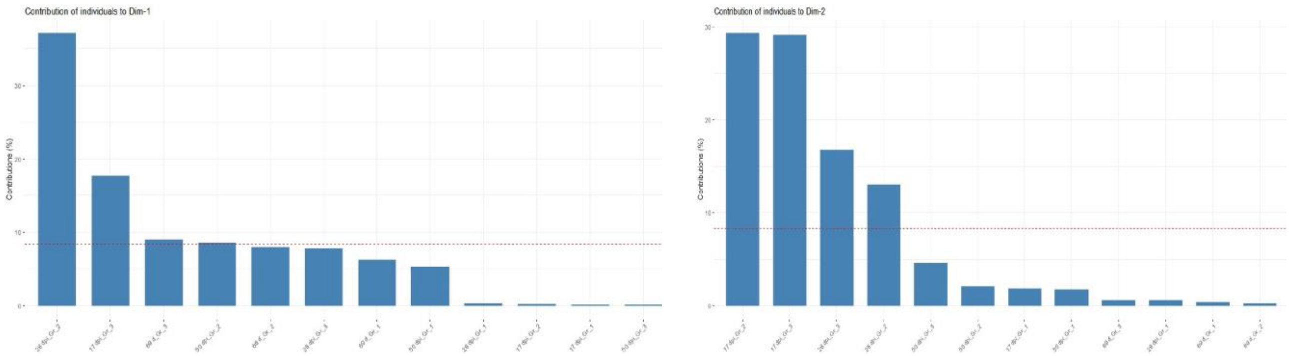


D

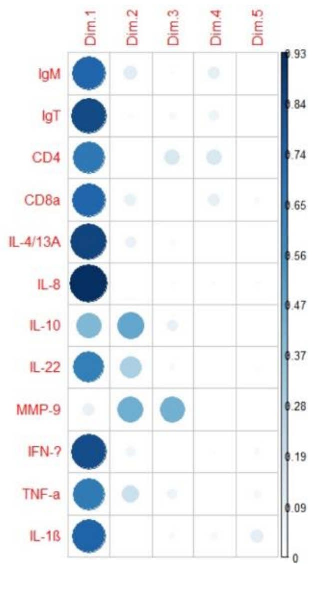


Significant gene variables in PC1

| | correlation | p.value |
|---|---|---|
| IL-8 | 0.9649025 | 3.954169e-07 |
| IL-4/13A | 0.9248497 | 1.662333e-05 |
| IgT | 0.9153853 | 2.959569e-05 |
| IFN-γ | 0.9069430 | 4.692682e-05 |
| IL-1ß | 0.8669219 | 2.617845e-04 |
| CD8α | 0.8620786 | 3.103661e-04 |
| IgM | 0.8609061 | 3.231173e-04 |
| CD4 | 0.8135278 | 1.286323e-03 |
| TNF-α | 0.8092930 | 1.428278e-03 |
| IL-22 | 0.8003650 | 1.766801e-03 |
| IL-10 | 0.6456387 | 2.334714e-02 |

Significant gene variables in PC2

| | correlation | p.value |
|---|---|---|
| IL-10 | 0.6914753 | 0.01274429 |
| MMP-9 | 0.6724146 | 0.01658941 |

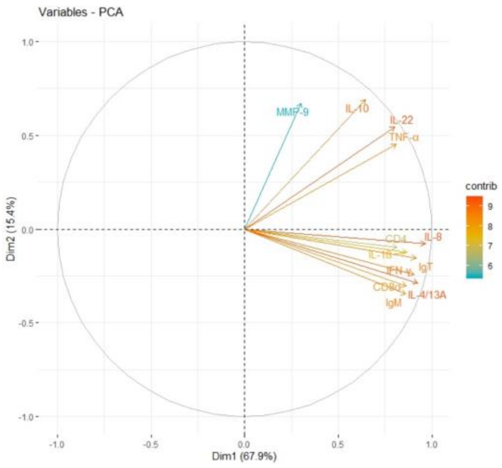
