## Supplementary figures for "Impact of a candidate vaccine on the dynamics of salmon lice (*Lepeophtheirus salmonis*) infestation and immune response in Atlantic salmon (*Salmo salar* L.)": S3_Fig.pdf

### PCA analysis- Spleen

A

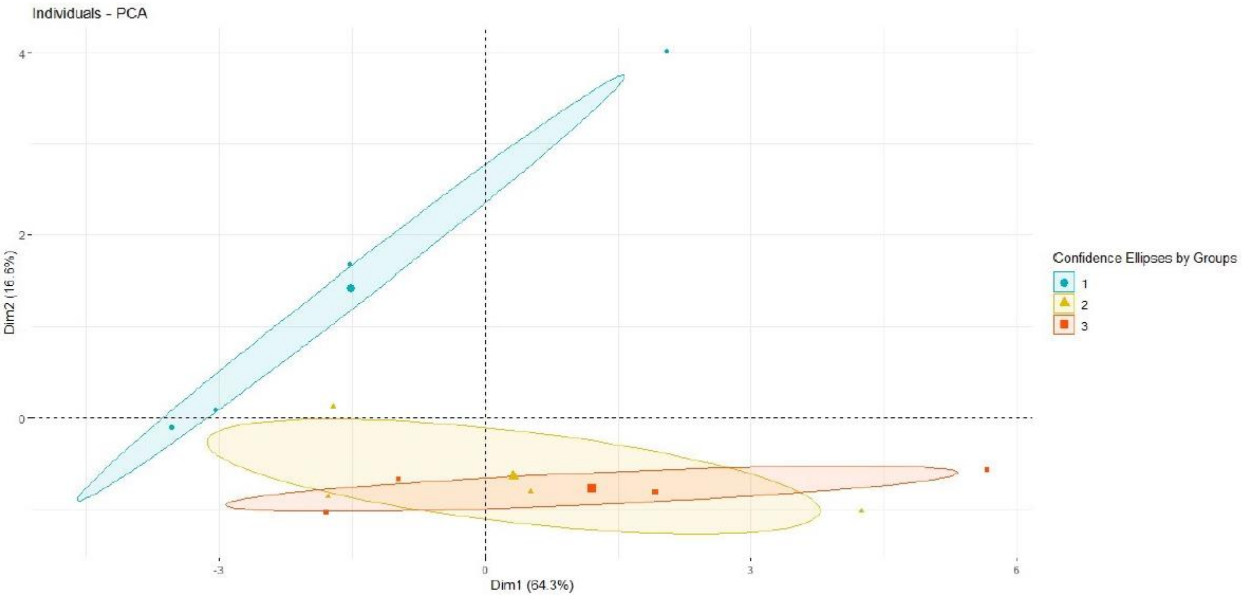


B

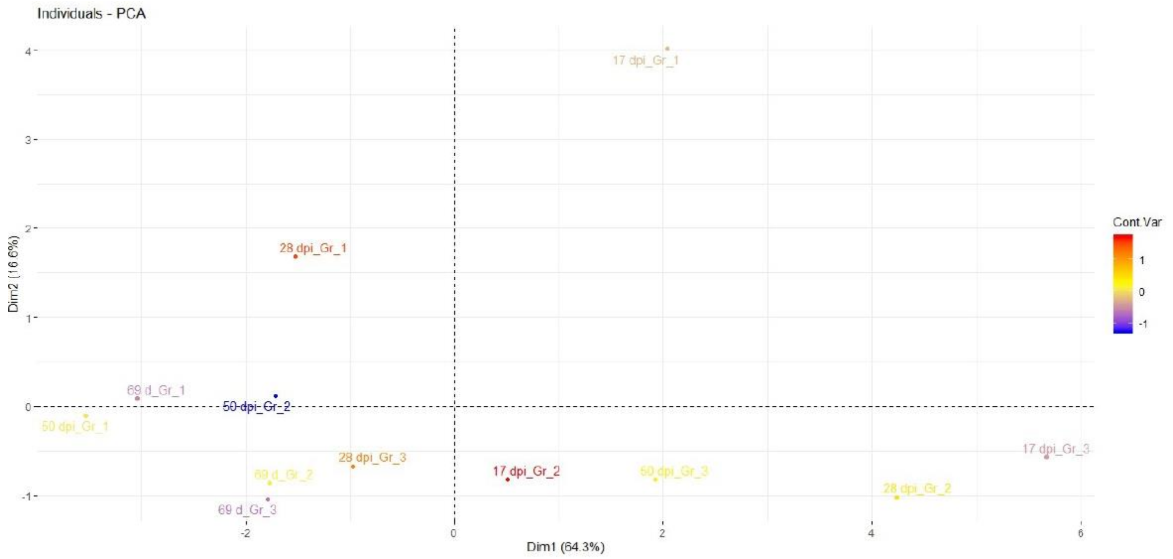


C

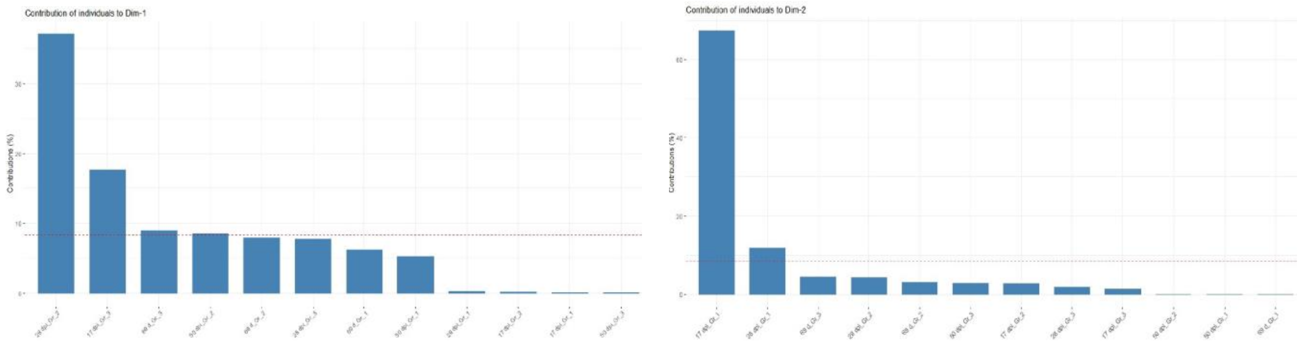


D

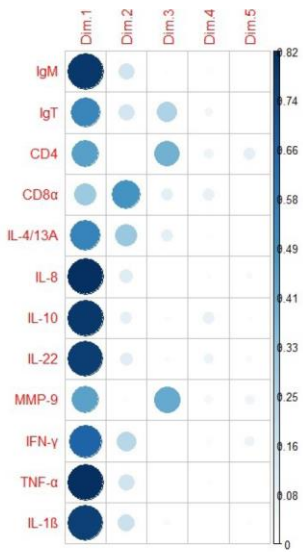


Significant gene variables in PC1

| | correlation | p.value |
|---|---|---|
| IL-8 | 0.9070025 | 4.678179e-05 |
| TNF-α | 0.9027424 | 5.809485e-05 |
| IgM | 0.8930517 | 9.185290e-05 |
| IL-10 | 0.8887951 | 1.108230e-04 |
| IL-22 | 0.8806855 | 1.553596e-04 |
| IL-1ß | 0.8748690 | 1.951115e-04 |
| IFN-γ | 0.8148761 | 1.243487e-03 |
| IL-4/13A | 0.7423462 | 5.691674e-03 |
| IgT | 0.7337079 | 6.605637e-03 |
| CD4 | 0.6670357 | 1.781376e-02 |
| MMP-9 | 0.6665850 | 1.791927e-02 |

Significant gene variables in PC2

| | correlation | p.value |
|---|---|---|
| CD8α | 0.7016583 | 0.01098374 |

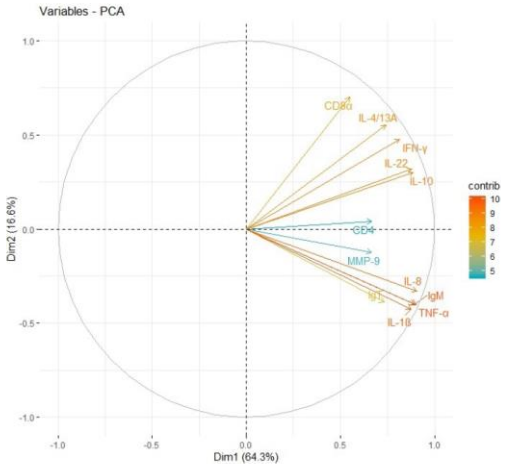
