## Supplementary figures and images for "Impact of a candidate vaccine on the dynamics of salmon lice (*Lepeophtheirus salmonis*) infestation and immune response in Atlantic salmon (*Salmo salar* L.)"

### S1_Fig.pdf

**Skin**

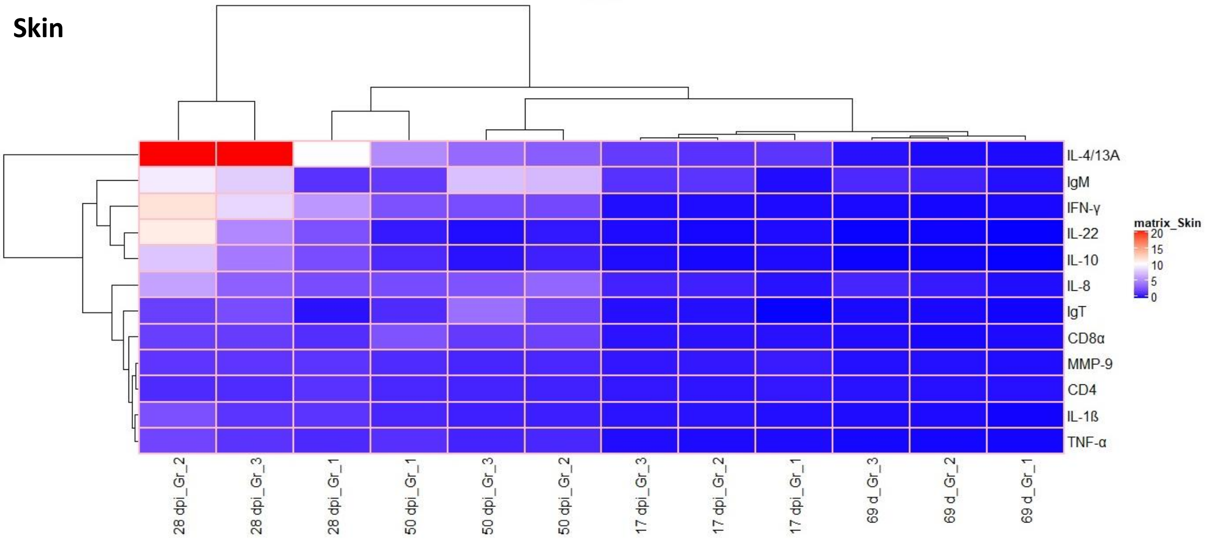


**Head kidney**

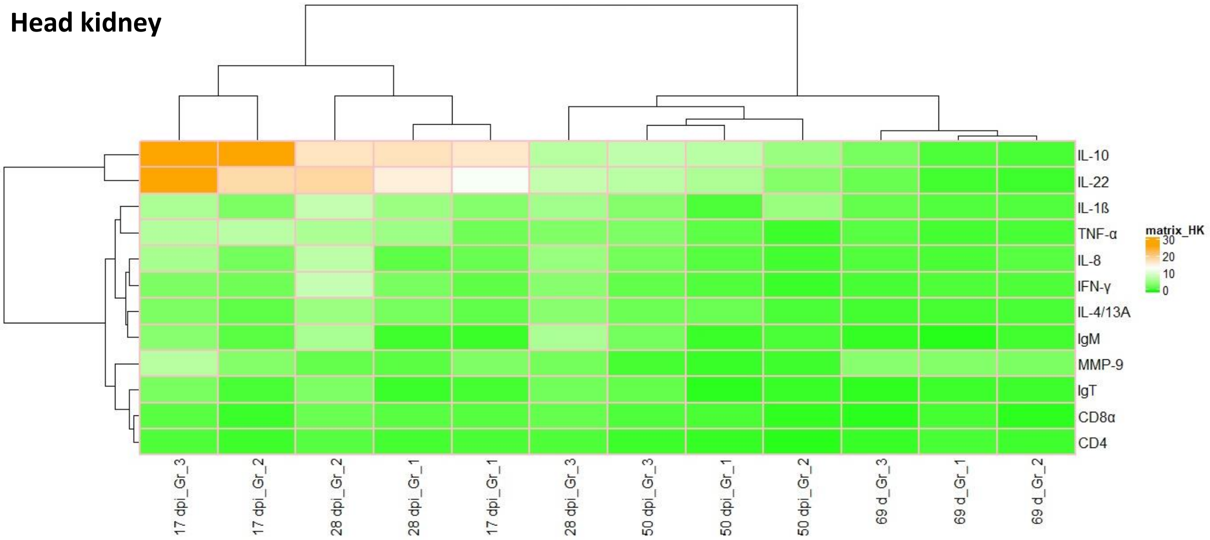


**Spleen**

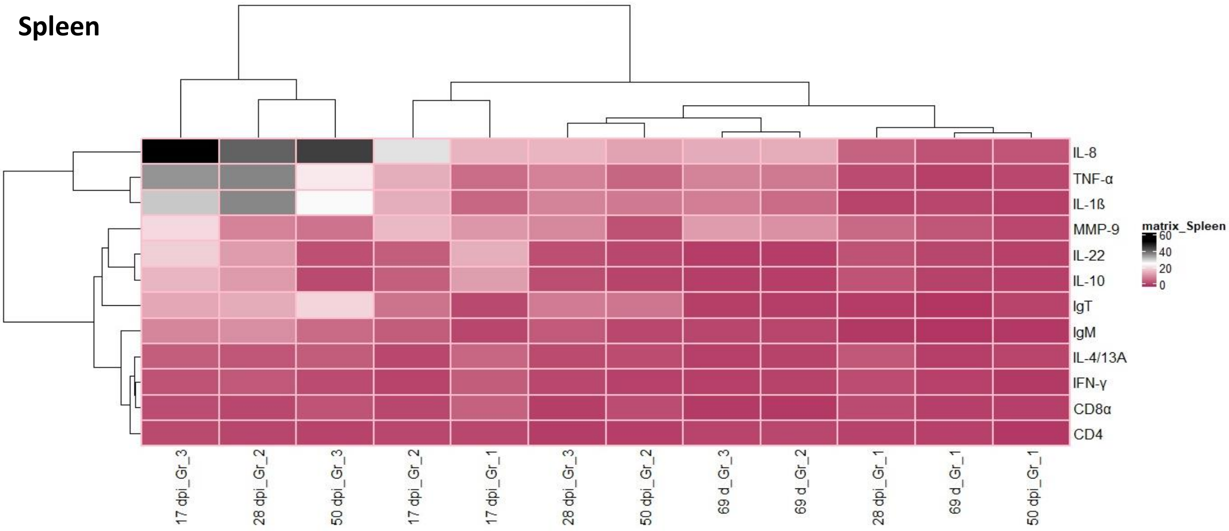

### S4-Fig.pdf

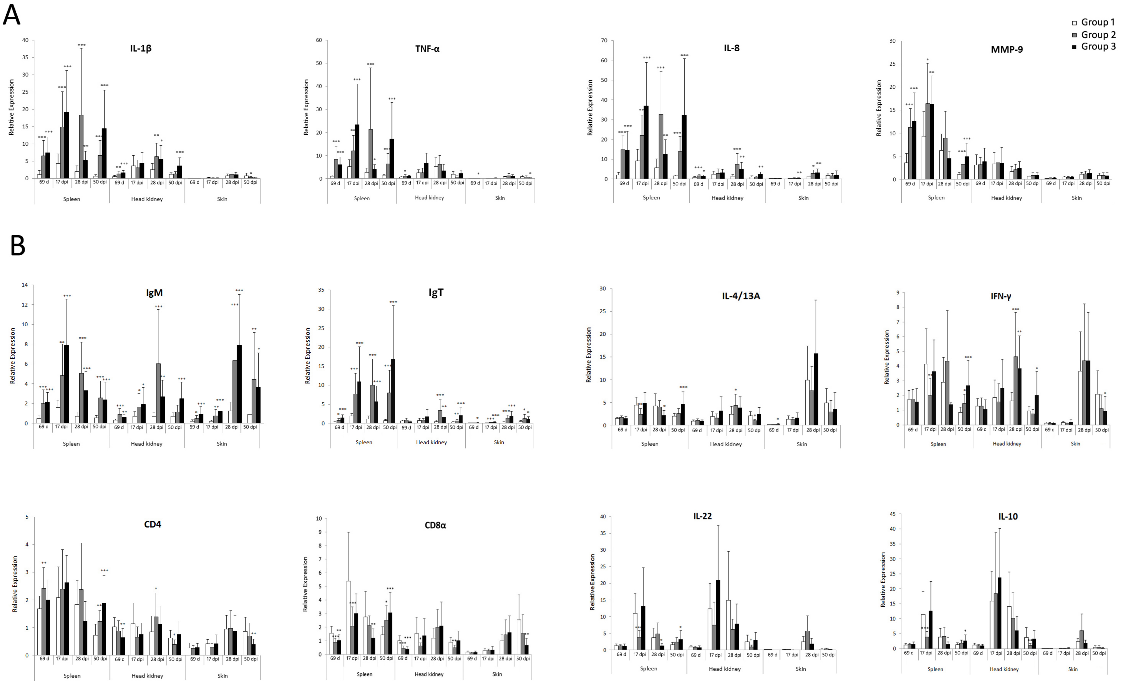
